## Supplementary Information for "Memory persistence and differentiation into antibody-secreting cells accompanied by positive selection in longitudinal BCR repertoires"

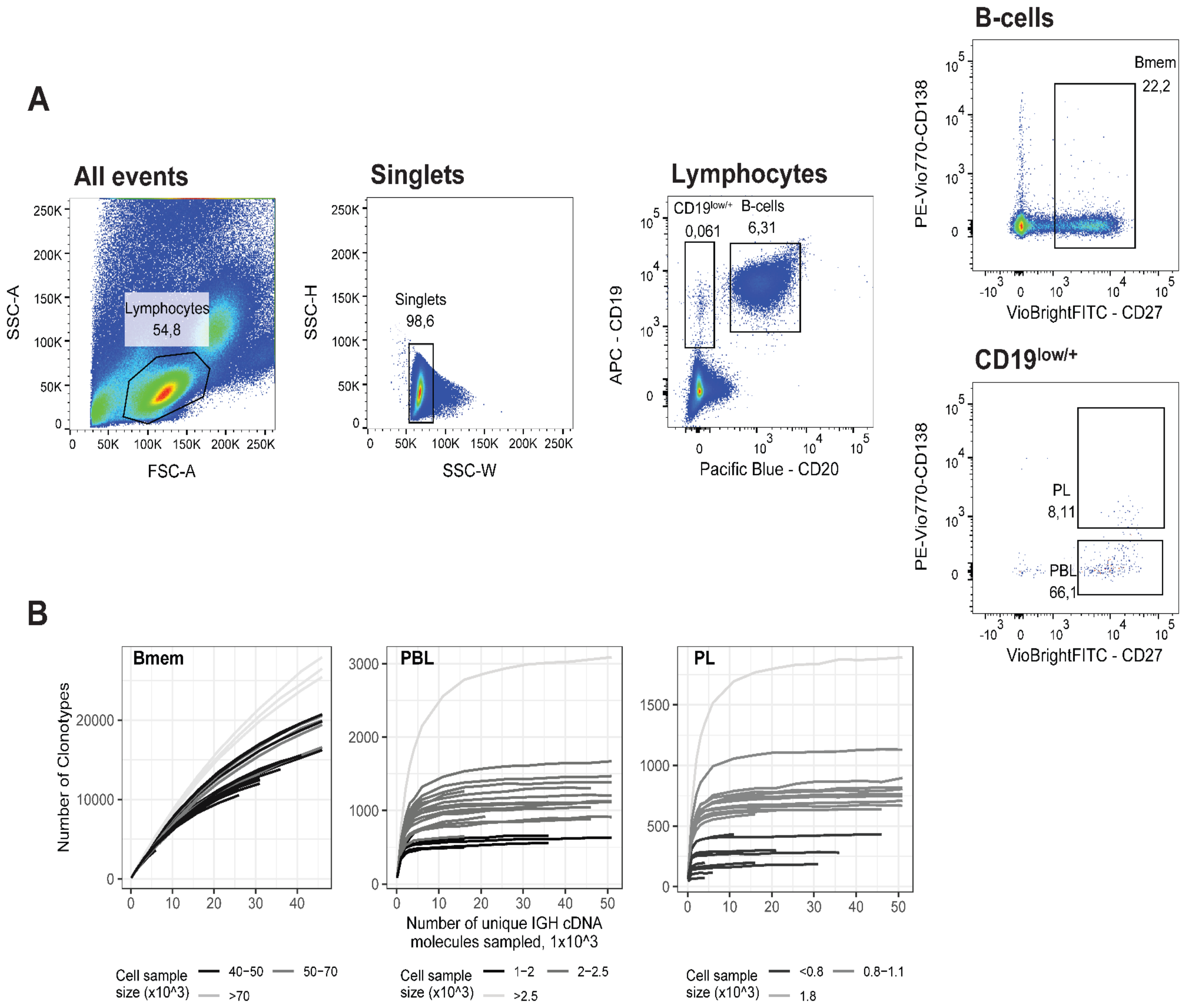


**Figure S1. A**: FACS gating strategy and frequencies of the following studied cell subsets for a representative peripheral blood sample (donor IZ, time point T3): Memory B-cells (Bmem: CD19$^{+}$ CD20$^{+}$ CD27$^{+}$), plasmablasts (PBL: CD19$^{low/+}$ CD20$^{-}$ CD27$^{high}$ CD138$^{-}$), and plasma cells (PL: CD19$^{low/+}$ CD20$^{-}$ CD27$^{high}$ CD138$^{+}$); **B**: Rarefaction curves for IGH cDNA molecules. From each repertoire, we sampled a defined number of unique IGH cDNA molecules and determined the number of unique IGH clonotypes. Each line represents a single sample. Shading of the lines indicates the number of cells sampled for each curve.

**Supplementary Table S1. Donor demographics and cell sample sizes.** Multiple values in a cell separated by a semicolon represent replicates collected for the corresponding donor, time point, or cellular subset. AR - allergic rhinitis; FA - food allergy; HD - healthy donor.

| | | | | Number of cells per sample | | | | | | | | |
|---|---|---|---|---|---|---|---|---|---|---|---|---|
| | | Time point | | | T1 | | | T2 | | | T3 | |
| Donor ID | Age | Sex | Status | Bmem | PBL | PL | Bmem | PBL | PL | Bmem | PBL | PL |
| D01 | 27 | F | AR | n/a | n/a | n/a | 50,300; 55,400 | 2,100; 2,100 | 1,020; 1,010 | 50,000; 50,000 | 1,000; 1,000 | 500; 500 |
| IM | 39 | M | AR,FA | 186,572 | 2,200 | 129 | 69,900; 68,400 | 2,000; 2,486 | 920 | 50,000; 50,000 | 2,000; 2,000 | 1,000; 1,000 |
| MRK | 27 | M | AR | 143,162 | 5,336 | 251 | 51,700; 50,600 | 2,130; 2,020 | 1,000; 1,035 | 50,000; 50,000 | 1,000; 1,000 | 400; 200 |
| AT | 23 | M | AR,FA | 101,400 | 7,200 | 1,800 | 50,600; 57,400 | 2,520 | 800 | 50000; 40800 | 1,000; 1,000 | 400; 200 |
| IZ | 33 | M | HD | 101,800 | 3,900 | 850 | 50,500; 56,300 | 1,140; 1,840 | 1,050; 625 | 50,000; 50,000 | 2,000; 2,000 | 200; 200 |
| MT | 33 | F | HD | n/a | n/a | n/a | n/a | n/a | n/a | 50,000; 50,000 | 1,000; 1,000 | 400 |

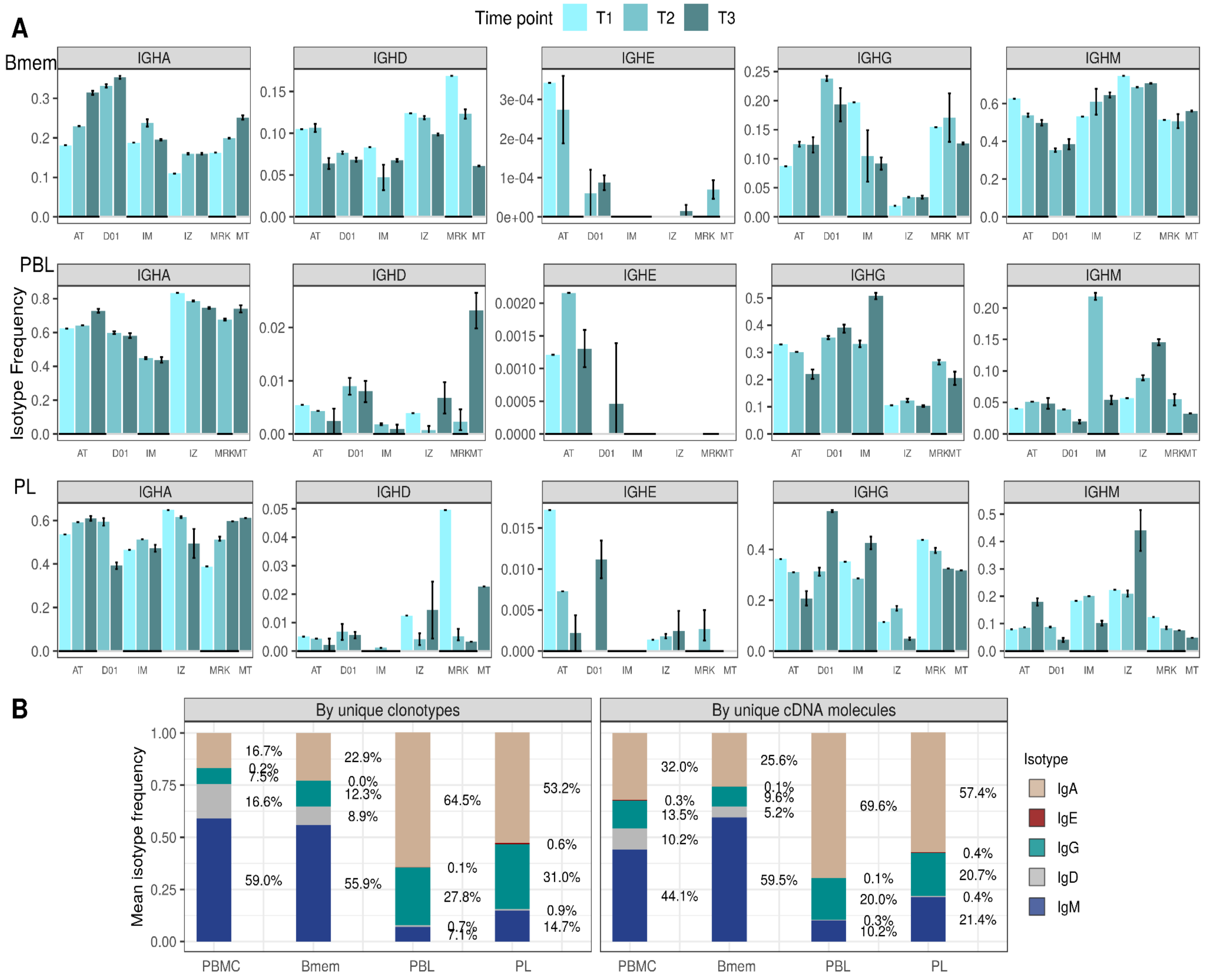


**Figure S2. Isotype frequencies in studied subsets. A:** Isotype frequencies in studied cell subsets and bulk PBMCs averaged across all samples. Frequencies were calculated as the number of IGH clonotypes (full-length unique nucleotide sequence), with specific isotypes divided by total number of clonotypes (left), or as the number of cDNA molecules in each isotype divided by the total number of cDNA molecules. **B**: Isotype frequencies based on unique clonotypes for different cell types in each individual donor sample. Whiskers illustrate minimal and maximal isotype frequencies for the group. Black and gray lines at the bottom of the plot indicate groups of bars corresponding to a particular donor.

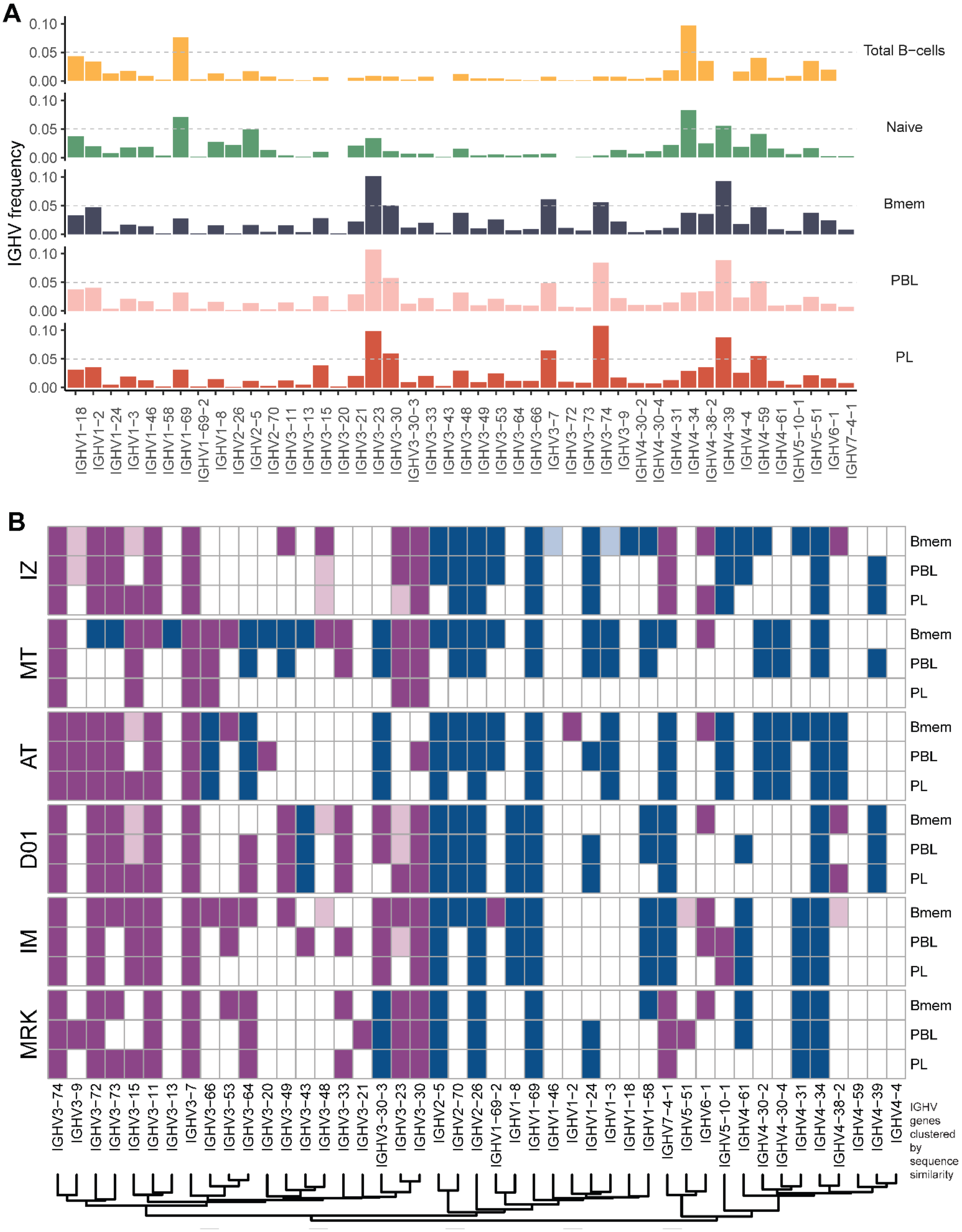
A
B
IGHV frequency
Total B−cells
Naive
Bmem
PBL
PL
IZ
MT
AT
D01
IM
MRK
IGHV genes clustered by sequence similarity

**Figure S3. IGHV gene frequencies in studied cell subsets. A**: Distributions of average IGHV gene frequencies in repertoires of total B cells, naive B cells (from Gidoni et al. 2019), Bmem, PBL, and PL cells. **B**: Heatmaps of IGHV frequencies for individual donors. Colored squares on heatmap indicate significantly different (FDR < 0.01) IGHV gene segment usage frequency in corresponding B cell subsets versus publicly available naive B cell repertoires. Color intensity reflects magnitude difference (FC = fold change). Only V genes represented by more than two clonotypes on average are shown. IGHV gene segments are ordered by similarity of their amino acid sequence, as indicated by the dendrogram at the bottom.

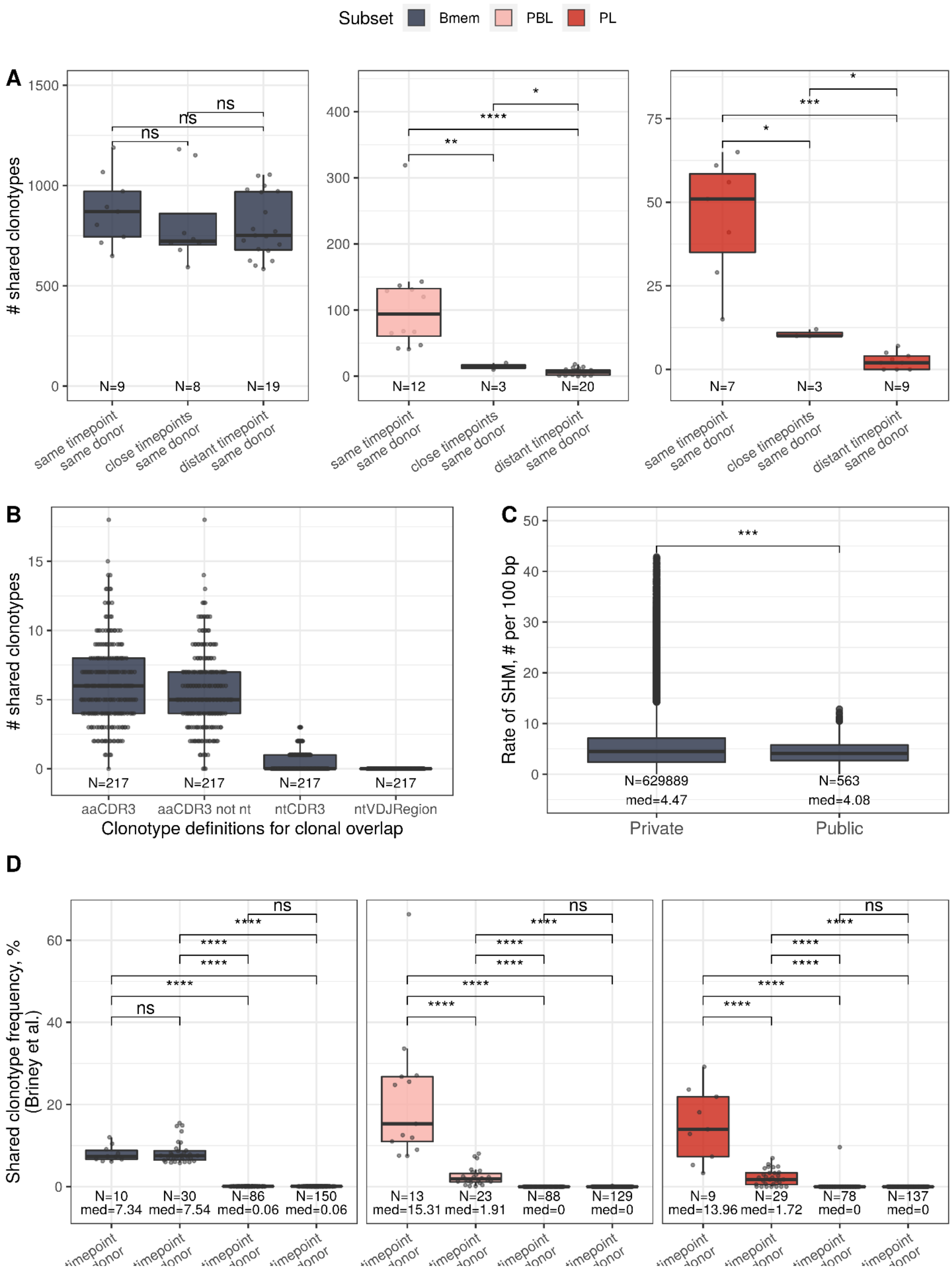
Subset
Bmem
PBL
PL
A
### shared clonotypes
N=9
N=8
N=19
N=12
N=3
N=20
N=7
N=3
N=9
ns
same timepoint same donor
close timepoints same donor
distant timepoint same donor
B
### shared clonotypes
N=217
aaCDR3
aaCDR3 not nt
ntCDR3
ntVDJRegion
Clonotype definitions for clonal overlap
C
Rate of SHM, # per 100 bp
N=629889 med=4.47
N=563 med=4.08
Private
Public
D
Shared clonotype frequency, % (Briney et al.)
N=10 med=7.34
N=30 med=7.54
N=86 med=0.06
N=150 med=0.06
N=13 med=15.31
N=23 med=1.91
N=88 med=0
N=129 med=0
N=9 med=13.96
N=29 med=1.72
N=78 med=0
N=137 med=0

**Figure S4. IGH repertoire similarity within B cell lineage subpopulations. A**: Number of shared clonotypes from each B cell subset between repertoires from a given donor at various time-points. “Same” represents replicate samples from the same blood draw, “close” samples were collected within ~1 month interval of each other, and “distant” samples were separated by an ~11–12 month interval. **B**: Number of shared clonotypes between pairs of repertoires from different donors with different clonotype definitions used to calculate overlap: aaCDR3 = amino acid CDR3 sequence and V gene label; aaCDR3 not nt = amino acid CDR3 sequence and V gene label, excluding clonotypes with identical CDR3 nucleotide sequence; ntCDR3 = nucleotide CDR3 sequence and V gene label; ntVDJRegion = full nucleotide sequence from the beginning of the IGH Framework 1 region to the end of the IGH Framework 4 region. **C**: Distribution of the number of somatic hypermutations identified per 100 bp length of IGHV segment for clonotypes detected either in only one donor (private) or in at least two donors (public). **D**: Shared clonotype frequency between pairs of repertoires, calculated as in Briney et al. 2019. For normalization purposes, we considered the 14,000 most abundant Bmem clonotypes, the 600 most abundant PBL clonotypes, and the 300 most abundant PL clonotypes. Dots in each plot represent pairs of repertoires of corresponding type. N = the number of pairs of repertoires in the group, and med = the median value. Comparisons in all panels were performed with Mann-Whitney test. * = $p \leq 0.05$, ** = $p \leq 0.01$, *** = $p \leq 10^{-3}$, **** = $p \leq 10^{-4}$.

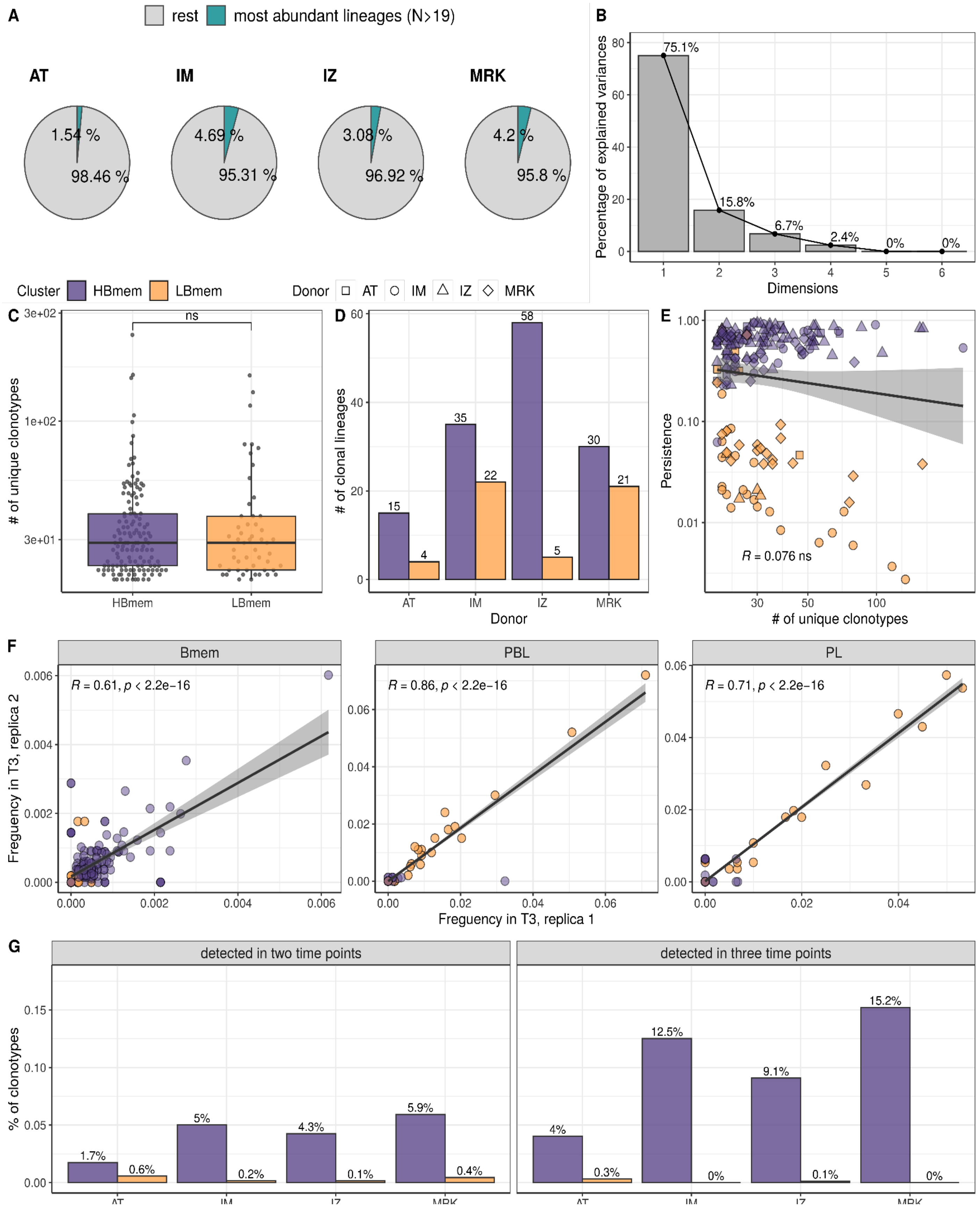

A
rest
most abundant lineages (N>19)
AT
IM
IZ
MRK
1.54 %
98.46 %
4.69 %
95.31 %
3.08 %
96.92 %
4.2 %
95.8 %
B
Percentage of explained variances
Dimensions
75.1%
15.8%
6.7%
2.4%
0%
0%
Cluster
HBmem
LBmem
Donor
AT
IM
IZ
MRK
C
### of unique clonotypes
ns
HBmem
LBmem
D
### of clonal lineages
Donor
15
4
35
22
58
5
30
21
E
Persistence
### of unique clonotypes
R = 0.076 ns
F
Bmem
PBL
PL
R = 0.61, p < 2.2e−16
R = 0.86, p < 2.2e−16
R = 0.71, p < 2.2e−16
Frequency in T3, replica 2
Frequency in T3, replica 1
G
detected in two time points
detected in three time points
% of clonotypes
1.7%
0.6%
5%
0.2%
4.3%
0.1%
5.9%
0.4%
4%
0.3%
12.5%
0%
9.1%
0.1%
15.2%
0%

**Figure S5. A**: Proportion of IGH clonotype diversity occupied by the most abundant clonal lineages (> 19 unique clonotypes). **B**: Scree plot for the principal component analysis (PCA) from **Fig. 3A** of the composition of clonal lineages, where fractions of Bmem, PBL, PL, and fractions of IgM, IgG and IgA were used as variables. **C**: Distribution of the number of unique clonotypes in a lineage for HBmem and LBmem. **D**: The number of clonal lineages belonging to HBmem or LBmem clusters in each donor. **E**: Spearman's correlation between the size of a clonal lineage and its persistence. **F**: Spearman's correlation between frequencies of clonal lineages in two replicates of time-point 3 (T3) samples. Only clonal lineages sampled with at least one replica at this time-point were included in the analysis; **G**: Fraction of clonotypes in HBmem or LBmem clonal lineages detected at two or three time-points.

**Supplementary Table S2.** Examples of divergent (*D*) and polymorphic (*P*) sites as calculated for the McDonald-Kreitman (MK) test. Synonymous and nonsynonymous substitutions from germ-line sequence are underlined and boldfaced, correspondingly.

| **Codon #** | **i** | | | **j** | | | **q** | | | **r** | | |
|---|---|---|---|---|---|---|---|---|---|---|---|---|
| **Germline** | c | g | c | g | - | - | c | t | a | a | a | t |
| **MRCA** | c | g | t | g | t | a | c | t | c | a | **g** | t |
| **Clonotypes in the clonal lineage** | c<br>c<br>c<br>c | g<br>**t**<br>g<br>**a** | t<br>t<br>t<br>t | g<br>g<br>g<br>g | t<br>t<br>t<br>t | a<br>a<br>a<br>a | c<br>c<br>c<br>c | t<br>t<br>t<br>t | c<br>a<br>c<br>c | a<br>a<br>a<br>a | **g**<br>**g**<br>**c**<br>**c** | t<br>t<br>t<br>t |
| ***Nonsynonymous divergence (Dn)*** | 0 | 0 | 0 | - | - | - | 0 | 0 | 0 | 0 | **1** | 0 |
| ***Synonymous divergence (Ds)*** | 0 | 0 | 1 | - | - | - | 0 | 0 | 0 | 0 | 0 | 0 |
| ***Nonsynonymous polymorphism (Pn)*** | 0 | **2** | 0 | - | - | - | 0 | 0 | 0 | 0 | **1** | 0 |
| ***Synonymous polymorphism (Ps)*** | 0 | 0 | 0 | - | - | - | 0 | 0 | 1 | 0 | 0 | 0 |
| **Comment** | example of codon represented by multiple variants in the clonal lineage (*i.e.* with the multiallelic site) | | | example of codon excluded from analysis because of unknown germline sequence for the site | | | example of codon where divergence is not counted because of presence of the germline variant among sequence variants in the lineage | | | example of codon where divergence is counted because there are no clonotypes identical to the germline sequence | | |

**Supplementary Table S3.** MK test results under different inclusion criterion for clonal lineages from HBmem and LBmem clusters, which makes it possible to deal with zero values in G-MRCA nonsynonymous or synonymous divergence. The LBmem cluster demonstrated consistent results of the MK test under all inclusion criteria, and the α of joined inside cluster divergence (combined SHM for all lineages belonging to the cluster) corresponds well to the median α among clonal lineages. The HBmem cluster is better suited for this type of filter, since it generally has much lower G-MRCA distance, and some clonal lineages have no divergence in MRCA from reconstructed portions of the germline sequence. Estimated α on joined cluster divergence in the HBmem cluster varies depending on the type of the filter employed, but is always lower than the α of LBmem. Additionally, consideration of all clonal lineages with addition of pseudocounts to *Dn* and *Ds* produces a negative median α, because the α of a clonal lineage with zero G-MRCA distance will always produce a negative α.

| **Inclusion criteria** | All clonal lineages. Pseudocounts are added to *Dn* and *Ds* to deal with zero values in the MK test of distinct clonal lineages. | | Clonal lineages with nonzero G-MRCA distance (at least one nonsynonymous or synonymous substitution). Pseudocounts are added to *Dn* and *Ds* to deal with zero values in the MK test of distinct clonal lineages. | | Clonal lineages with at least one nonsynonymous and synonymous substitution. No pseudocounts in *Dn* and *Ds* are required. | |
|---|---|---|---|---|---|---|
| **Cluster** | HBmem | LBmem | HBmem | LBmem | HBmem | LBmem |
| **# of filtered clonal lineages** | 138 | 52 | 68 | 49 | 18 | 29 |
| **Median α** | -0.46 | 0.55 | 0.18 | 0.57 | - 0.07 | 0.54 |
| **Mann-Whitney test** | $p = 2.9 \cdot 10^{-11}$ | | $p = 4.8 \cdot 10^{-6}$ | | p = 0.0028 | |
| **MK test on joined diversity of the cluster** | α = 0.58<br>$p = 4.97 \cdot 10^{-7}$ | α = 0.65<br>$p < 2.2 \cdot 10^{-16}$ | α = 0.61<br>$p = 6.05 \cdot 10^{-8}$ | α = 0.66<br>$p < 2.2 \cdot 10^{-16}$ | α = 0.26<br>p = 0.1004 | α = 0.56<br>$p = 2.05 \cdot 10^{-10}$ |
